# The wasting-associated metabolite succinate disrupts myogenesis and impairs skeletal muscle regeneration

**DOI:** 10.1101/822338

**Authors:** Paige C Arneson, Kelly A Hogan, Alexandra M Shin, Adrienne Samani, Aminah Jatoi, Jason D Doles

**Author notes:** Corresponding Author: Jason D Doles, Department of Biochemistry and Molecular Biology, Mayo Clinic, 200 First St SW, Guggenheim 16-11A1, Rochester, MN 55905.

## Abstract

**Background:** Muscle wasting is a debilitating co-morbidity affecting most advanced cancer patients. Alongside enhanced muscle catabolism, defects in muscle repair/regeneration contribute to cancer-associated wasting. Among the factors implicated in suppression of muscle regeneration are cytokines that interfere with myogenic signal transduction pathways. Less understood is how other cancer/wasting-associated cues, such as metabolites, contribute to muscle dysfunction. This study investigates how the metabolite succinate affects myogenesis and muscle regeneration.

**Methods:** We leveraged an established ectopic metabolite treatment (cell permeable dimethyl-succinate) strategy to evaluate the ability of intracellular succinate elevation to 1) affect myoblast homeostasis (proliferation, apoptosis), 2) disrupt protein dynamics and induce wasting-associated atrophy, and 3) modulate *in vitro* myogenesis. *In vivo* succinate supplementation experiments (2% succinate, 1% sucrose vehicle) were used to corroborate and extend *in vitro* observations. Metabolic profiling and functional metabolic studies were then performed to investigate the impact of succinate elevation on mitochondria function.

**Results:** We found that *in vitro* succinate supplementation elevated intracellular succinate about 2-fold, and did not have an impact on proliferation or apoptosis of C2C12 myoblasts. Elevated succinate had minor effects on protein homeostasis (∼25% decrease in protein synthesis assessed by OPP staining), and no significant effect on myotube atrophy. Succinate elevation interfered with *in vitro* myoblast differentiation, characterized by significant decreases in late markers of myogenesis and fewer nuclei per myosin heavy chain positive structure (assessed by immunofluorescence staining). While mice orally administered succinate did not exhibit changes in overall body composition or whole muscle weights, these mice displayed smaller muscle myofiber diameters (∼6% decrease in the mean of non-linear regression curves fit to the histograms of minimum feret diameter distribution), which was exacerbated when muscle regeneration was induced with barium chloride injury. Significant decreases in the mean of non-linear regression curves fit to the histograms of minimum feret diameter distributions were observed 7 days and 28 days post injury. Elevated numbers of myogenin positive cells (3-fold increase) supportive of the differentiation defects observed *in vitro* were observed 28 days post injury. Metabolic profiling and functional metabolic assessment of myoblasts revealed that succinate elevation caused both widespread metabolic changes and significantly lowered maximal cellular respiration (∼35% decrease).

**Conclusions:** This study broadens the repertoire of wasting-associated factors that can directly modulate muscle progenitor cell function and strengthens the hypothesis that metabolic derangements are significant contributors to impaired muscle regeneration, an important aspect of cancer-associated muscle wasting.

## INTRODUCTION

Lean mass (muscle) loss and functional skeletal muscle decline affect the majority of advanced cancer patients and are associated with poor prognosis and quality of life. Reversing muscle loss in tumor-bearing animal models leads to improved chemotherapeutic efficacy, reduced cancer-associated morbidity, and prolonged survival^1,2^, demonstrating that maintaining muscle mass in cancer patients should be a top clinical priority. Despite a growing number of active clinical trials for cancer-associated weight loss, effective therapies remain elusive. Major problems hampering significant progress include the variability of weight loss presentation across tumor types and stages, limited early detection of lean mass/adipose tissue loss, and a poor understanding of the molecular underpinnings of this complex syndrome. More work is needed to identify and target alterations in skeletal muscle that contribute to overall tissue decline in cancer patients.

Multiple factors produced by both tumor and host contribute to the wasting state. Many identified ‘wasting factors’ are cytokines/chemokines such as TNF-alpha, Interleukins (including IL-1 and IL-6), CXC-proteins, and TGF-β^3-6^, and are known to accelerate muscle atrophy, promote protein breakdown, and inhibit muscle regeneration. Additionally, many of these factors are linked to mitochondria dysfunction and altered cellular metabolism^7^. Indeed, metabolic regulation of skeletal muscle function has come into focus as a major driver of multiple aspects of muscle biology including catabolism, hypertrophy, and regenerative capacity. Examples of metabolic processes linked to cancer-associated muscle wasting are: mitochondria dynamics^8^, mitochondria biogenesis^9,10^, mitochondria catabolism (ie. mitophagy)^11^, and compromised energy utilization pathways^12^. A more sophisticated understanding of how metabolic factors contribute to cancer-associated muscle dysfunction is needed to facilitate the development of novel therapies to preserve muscle mass and function in cancer patients.

Succinate is a metabolite best known for its role as a TCA cycle intermediate and central player in mitochondria metabolism and ATP generation. Recently, new roles for succinate have emerged and include immune cell modulation^13^, HIF-1a stabilization in cancer^14^, epigenome remodeling via inhibition of alpha-ketoglutarate dependent dioxygenases^15^, and post-translational protein modification (succinylation)^16^. This expanding repertoire of cellular activities suggests a broader role for succinate in regulating cellular processes such as differentiation, fate decisions, and survival. Interestingly, a recent study reported serum succinate accumulation as a biomarker capable of distinguishing non-wasting, tumor-bearing, and tumor-bearing with weight loss patients^17^. In the present study, we sought to determine if succinate elevation directly impacts muscle cell function, with a particular emphasis on myogenic differentiation given prior studies linking succinate levels to cell fate decisions^18,19^.

Alterations in the balance between muscle breakdown and repair/regeneration underlie cancer-associated muscle wasting. While catabolic pathways have historically received the most attention in the muscle wasting field, processes leading to the suppression of muscle regeneration are understudied despite their clear contribution to the etiology of muscle wasting^20^. Central to the process of muscle regeneration is the skeletal muscle stem cell, or satellite cell. In healthy muscle, satellite cells contribute to tissue maintenance and actively participate in muscle homeostasis^21,22^. When challenged by injury stimuli, satellite cells activate, proliferate, and differentiate into mature muscle cells in order to facilitate muscle repair and regeneration. Indeed, genetic ablation experiments show that Pax7+ stem/satellite cells are required for muscle regeneration following major trauma^23-25^. Defects in satellite cell function and/or regeneration are associated with a growing list of muscle atrophy states including disuse^26,27^, denervation^28,29^, chronic obstructive pulmonary disease (COPD) ^30^, burn injury^31,32^, diabetes^33,34^, chronic kidney disease^35^, and hepatic disorders^36^, although the precise role of impaired repair/regeneration in each of these disease states remains unclear. In cancer, multiple studies show that muscle regeneration is linked to functional skeletal muscle deficits and propose that mononuclear muscle progenitor cells drive aspects of the atrophy phenotype^37,38^. Taking into account the expanding role of the satellite cell in muscle wasting, the importance of metabolism in cancer-associated wasting, and recent evidence implicating the metabolite succinate in cancer cachexia^17^ and stem/progenitor cell function^18,19^, we sought to characterize the effects of succinate elevation on muscle myoblast cell dynamics (*in vitro*) and muscle homeostasis (*in vivo*). Studies presented herein highlight the ability of a metabolite – succinate – to directly impact myogenesis and muscle regeneration. This underscores the need to further explore metabolic drivers of impaired muscle regeneration and muscle catabolism, both contributors to cancer-associated muscle wasting, in order to advance anti-wasting cancer therapies.

## METHODS

### Animals

Mice were bred and housed according to NIH guidelines for the ethical treatment of animals in a pathogen-free facility at the Mayo Clinic (Rochester, MN campus). The Mayo Clinic Institutional Animal Care and Use Committee (IACUC) approved all animal protocols and all experiments were performed in accordance with relevant guidelines and regulations. Wild type mice were on the FVB background (Charles River Labs) unless noted otherwise in the text or figure legends. For homeostasis and regeneration assays, all mice were 3-5 months old, FVB males. In these experiments, mice were administered succinate (Sigma-Aldrich) (2%) and 1% sucrose (vehicle) or 1% sucrose (vehicle only) in their drinking water for 6-8 weeks prior to and throughout the injury time course. Muscle injury was performed by injecting the left *tibialis anterior* muscle with 70uL BaCl_2_ (1.2% in saline) or saline (control) and harvesting muscles 1 or 4 weeks post injury.

### EchoMRI imaging

An EchoMRI (magnetic resonance imaging) Body Composition Analyzer (Echo Medical Systems, Houston, USA) was used for longitudinal body composition analyses. Mice were scanned twice weekly for the 6-week duration of the succinate water experiments. All body composition analyses were performed in the accompanying EchoMRI software. Mice were weighed on a standard, digital lab scale prior to scanning.

### Cell culture

C2C12 cells (ATCC CRL-1772) were grown on tissue culture treated dishes in growth media consisting of high glucose DMEM media supplemented with 10% fetal bovine serum and 1% penicillin/streptomycin. Differentiation-inducing media contains 2% horse serum in place of 10% fetal bovine serum. Differentiating cultures were maintained by alternating fresh media additions and complete media changes on a daily basis for 5 days. DM-succinate (Sigma-Aldrich) was added directly to the culture media upon switching to differentiation media at doses of 4 or 8 mM. <u>Live cell analyses</u>: Cell proliferation and cell death were measured by live cell analysis (Incucyte S3 Live-Cell Imaging System, Essen Bioscience). Percent apoptotic cells over a 48h time course was determined by labeling control or treated cultured C2C12 cells with a phosphatidylserine cyanine fluorescent dye (Annexin V Red Reagent, Essen Bioscience) according to the manufacturer.

### Immunostaining

Tissue for immunostaining was placed in a sucrose sink (30%) overnight prior to freezing and sectioning. Sections (8-10 um) were post-fixed in 4% paraformaldehyde (PFA) for 5 minutes at room temperature prior to immunostaining. Once fixed, tissue sections were permeabilized with 0.5% Triton-X100 in PBS followed by blocking with 3% BSA, 0.2% Triton-X and 0.2% Tween-20 in PBS. Primary antibody incubations occurred at RT for 90 minutes followed by incubation with secondary antibody at RT for 30 minutes in buffer described above. The following antibodies were used in this study: MF-20 (Developmental Hybridoma Bank), Laminin (Sigma 4HB-2), and BF-F3 (Developmental Hybridoma Bank), and SC-71 (Developmental Hybridoma Bank), Pax7 (Developmental Hybridoma Bank), Ki67 (Santa Cruz) and Myogenin (Santa Cruz). Secondary antibodies were all Alexa fluorescent conjugates (488, 555, or 647) from Invitrogen or Jackson ImmunoResearch.

### Flow cytometry

#### OPP assay

C2C12 myoblasts were grown to 60% confluency prior to treatment with either control media or 8 mM succinate media for 48 hours. N= 3 plates control, n=3 8mM succinate. Cells were incubated with O-Propargyl-Puromycin (OPP) for 1.5 hours and then trypsinized and stained following the manufacturer’s instructions (protein synthesis assay kit (Cayman Chemical #601100)). Viobility 405/452 fixable dye (Miltenyi Biotec 130-110-205) was used as live/dead discriminator. After staining, cells were immediately analyzed by flow cytometry on a MACSquant 10 analyzer (Miltenyi Biotec). Data were analyzed using MACSquantify software (Miltenyi Biotec).

### Quantitative RT-qPCR

C2C12 cells were differentiated as described above and cell pellets were collected at each day post differentiation as well as prior to differentiation (day 0, growth media). RNA was isolated using column purification (Qiagen) and cDNA was prepared using high capacity cDNA reverse transcription kit (Applied Biosystems). qPCR was preformed using BioRad CFX384 Real Time System. Primer sequences are available upon request.

### Metabolic Assays

A Seahorse XFe24 Analyzer (Agilent) was used to perform mitochondria stress tests and energy phenotyping tests on C2C12 cells. Cells were treated with 4 or 8mM DM-succinate or control growth media for 16-18 hours prior to the assay. DM-succinate treatment was maintained throughout the assay. Oligomycin (1 μM), FCCP (0.4 μM) and rotenone/antimycin A (1 μM, 1 μM) were used to inhibit mitochondria complexes involved in electron transport/ATP production. Prior to start of the assay, total cells/well counts were obtained on a Celigo imaging cytometer (Nexcelom Bioscience) using Hoechst 33342 stain (Thermofisher). All data were normalized to Celigo counts of cells/well. Wave software (Agilent, version 3.0.11) was used in the analysis of energy phenotyping data.

### Quantitative metabolite analyses

#### Sample preparation

Cells were treated for 48 hours with 4 or 8 mM DM-succinate, washed 3 times with PBS, and stored at −80C until metabolite extraction. Muscle tissue was cut into ∼50 mg pieces, flash frozen in liquid nitrogen, and stored at −80 C until metabolite extraction. Conditioned media was flash frozen in liquid nitrogen. All samples were submitted to the Mayo Clinic Metabolomics Resource Core for targeted metabolomics.

#### TCA

Concentration of TCA analytes were measured by gas chromatograph mass spectrometry (GC/MS) as previously described with a few modifications^39^. Briefly, cell culture supernatants were spiked with internal solutions containing U-^13^C labeled analytes. Proteins were removed by adding 250 ul of chilled methanol and acetonitrile solution to the sample mixture. After drying the supernatant in the speed vac, the sample was derivatized with ethoxime and then with MtBSTFA + 1% tBDMCS (N-Methyl-N-(t-Butyldimethylsilyl)-Trifluoroacetamide + 1% t-Butyldimethylchlorosilane) before it was analyzed on an Agilent 5975C GC/MS (gas chromatography/mass spectrometry) under electron impact and single ion monitoring conditions. Concentrations of lactic acid (m/z 261.2), fummaric acid (m/z 287.1), succinic acid (m/z 289.1), oxaloacetic acid (m/z 346.2), ketoglutaric acid (m/z 360.2), malic acid (m/z 419.3), aspartic acid (m/z 418.2), 2-hydroxyglutaratic acid (m/z 433.2), cis-aconitic acid (m/z459.3), citric acid (m/z 591.4), and isocitric acid (m/z 591.4), glutamic acid (m/z 432.4) were measured against a 7-point calibration curves that underwent the same derivatization. n=1 for each supernatant sample, which was run in 2 technical replicates after derivatization.

### Untargeted metabolomics analyses

Three biological replicates of control and DM-succinate (8 mM) treated C2C12 cells were prepared for untargeted metabolomics analyses as follows: cells were grown to 60% confluency and then treated with 8 mM succinate for 48 hours, at which time plates were washed with PBS and cells were scraped and pelleted, followed by storage at −80° C. Samples were submitted to the Metabolomics Core at Mayo Clinic (Rochester, MN, USA) for metabolomics profiling by liquid chromatography-mass spectrometry (LC/MS) using 6550 iFunnel Quadrupole Time of Flight (Q-TOF) mass spectrometer (Agilent). Metabolites were then identified using METLIN database with Mass Profiler Professional software (Agilent). Annotated metabolites with a p value less than 0.05 and a fold change greater than 2 were further analyzed using MetaboAnalyst network explorer^40^ (metabolite-metabolite interactions) and enrichment analysis (pathway associated metabolite sets) tools. KEGG and HMDB identifiers were used in MetaboAnalyst.

### RNA sequencing analysis

#### Sample/library preparation

Three biological replicates of control and DM-succinate (8mM) treated C2C12 cells were prepared for RNA sequencing analyses as follows: cells were grown to 60% confluency and then treated with 8mM succinate for 48 hours, at which time plates were washed with PBS and cells were scraped and pelleted, followed by storage at −80° C. Frozen cell pellets were submitted to the Mayo Clinic Medical Genome Facility where RNA quality was determined using the Fragment Analyzer from AATI. RNA samples that have RQN values ≥6 were approved for library prep and sequencing. RNA libraries were prepared using 200 ng of good quality total RNA according to the manufacturer’s instructions for the TruSeq RNA Sample Prep Kit v2 (Illumina, San Diego, CA), employing poly-A mRNA enrichment using oligo dT magnetic beads. The final adapter-modified cDNA fragments were enriched by 12 cycles of PCR using Illumina TruSeq PCR primers. The concentration and size distribution of the completed libraries were determined using a Fragment Analyzer (AATI, Ankeny, IA) and Qubit fluorometry (Invitrogen, Carlsbad, CA). Libraries were sequenced following Illumina’s standard protocol using the Illumina cBot and HiSeq 3000/4000 PE Cluster Kit, yielding approximately 48-70 million fragment reads per sample. The flow cells were sequenced as 100 × 2 paired end reads on an Illumina HiSeq 4000 using HiSeq 3000/4000 sequencing kit and HCS v3.3.52 collection software. Base-calling was performed using Illumina’s RTA version 2.7.3.<u>Bioinformatics/Data Processing</u>: All bioinformatics analyses were done through the Mayo Clinic Bioinformatics Core. The raw RNA sequencing paired-end reads for the samples were processed through the Mayo RNA-Seq bioinformatics pipeline, MAP-RSeq version 3.0.0^41^. Briefly, MAP-RSeq employs the splice-aware aligner, STAR^42^, to align reads to the reference mouse genome build mm10. Gene and exon expression quantification were performed using the Subread^43^ package to obtain both raw and normalized (RPKM – Reads Per Kilobase per Million mapped reads) reads.

### Statistics

Data are represented as the mean ± SD using GraphPad Prism unless noted otherwise in the figure legends. Quantification of nuclei per MyHC+ structure was analyzed using nonparametric, Mann-Whitney t-tests. Quantification of muscle cross sections using minimum feret diameter measurements was analyzed by non-linear regression (gaussian, least squares method) and compared between conditions using an extra sum-of-squares F test. All other comparisons between groups were performed using unpaired two-tailed student’s t tests or multiple t tests with Holm-Sidak multiple testing correction, as noted in figure legends. For all analyses, a p<0.05 was considered significant (denoted with *); a p<0.01 was denoted with ** unless noted otherwise in the figure legend.

### Data availability

The RNAseq dataset generated and analyzed during the current study is available in the Sequence Read Archive (SRA) (National Center for Biotechnology Information, NCBI), submission number <u>SUB6936750</u>, BioProject ID PRJNA605465. All other datasets generated during the current study are available from the corresponding author upon request.

## RESULTS

### Modeling succinate accumulation and evaluating consequences on growth and atrophy

To evaluate the effects of succinate on *in vitro* myoblast function, we first sought to adapt an established paradigm of intracellular succinate accumulation for use in myoblast cells. Succinate has very low cell permeability *in vitro*, so we began our studies using dimethyl-succinate (DM-succinate or DMS), a cell permeable succinate analog used in prior studies to elevate intracellular metabolite levels^18,19^. We treated proliferating C2C12 myoblasts for 48 hours with two concentrations of DM-succinate, 4 mM and 8 mM, and then performed targeted metabolomics to quantify levels of TCA cycle-associated metabolites. We found that 8 mM DM-succinate treatment resulted in a significant increase in intracellular succinate (∼2.5-fold) compared to control treatment, similar in magnitude to previously reported values^18^ and consistent with the degree of succinate accumulation in wasting cancer patients^17^. We did not detect statistically significant changes in most other TCA metabolites analyzed, with the exception of small (<2 fold) but significant increases in malate and glutamate (**Fig. 1A**). We chose to evaluate this dosing paradigm further to investigate the effects of intracellular succinate elevation on myogenesis. Cell proliferation and apoptosis are important regulators of myogenesis. To test if succinate would influence either of these processes, we evaluated cells treated with three doses of DM-succinate over 48 hours. We did not see significant differences in proliferation over this time period, and only saw increased apoptosis rates with 16 mM DM-succinate treatment (**Fig. 1B, C**).

**Figure 1:**
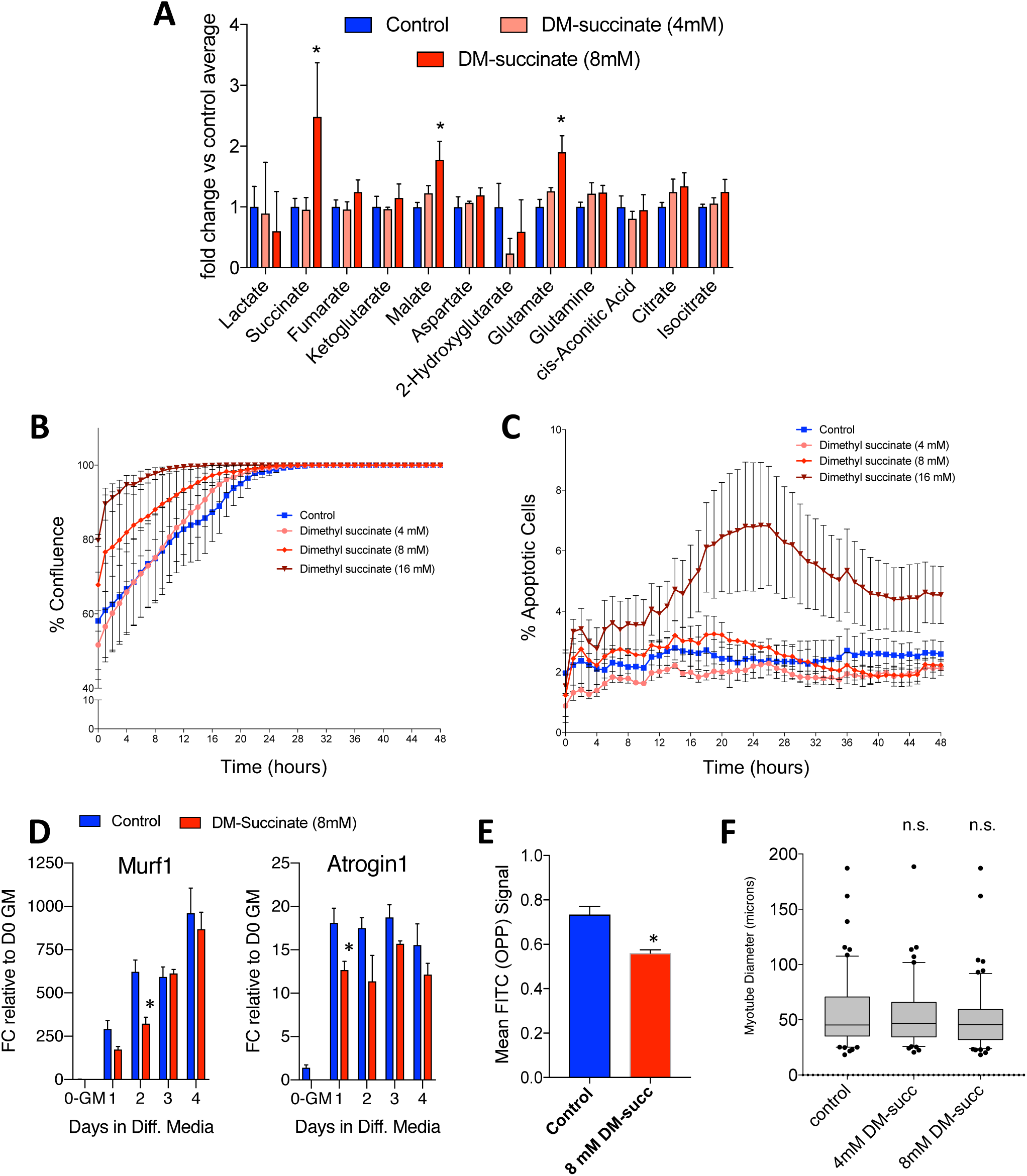
Modeling succinate accumulation and evaluating consequences on growth and atrophy. **(A)** A bar graph depicting relative intracellular levels of TCA metabolites in C2C12 myoblasts following 48h DM-succinate treatment as determined by gas chromatography mass spectrometry (GC/MS) analyses. *p<0.05. n=3 replicates. **(B)** Proliferation curve (measured as percent confluence) for control, 4, 8, and 16 mM DM-succinate over 48 hours. Percent confluence was averaged from 3 images per replicate, N= 6 replicate wells for each condition. Error bars represent SEM. **(C)** Percent apoptotic cells, measured by number of Annexin V positive cells, for control, 4, 8, and 16 mM DM-succinate over 48 hours. Percent apoptotic cells was averaged from 3 images per replicate, N=3 replicate wells for each condition. Error bars represent SD. **(D)** Bar graphs quantifying Murf1 (left) and Atrogin1 (right) mRNA expression FC relative to D0-growth media at 0, 1, 2, 3, and 4d in differentiation media +/- 8 mM DM-succinate. *p<0.05 between control and succinate samples for marked timepoints, as determined by multiple t tests with Holm-Sidak multiple testing correction. **(E)** A bar graph showing reduced OPP accumulation in C2C12 cells treated for 48h with 8mM DM-succinate. n=3 replicates; *p<0.05. **(F)** A graph of myotube diameter measurements (microns) from C2C12 myotubes treated with DM-succinate for 24h. n.s.=not significant (p>0.05) between any conditions tested.

A major feature of cancer-associated muscle wasting is myofiber atrophy due to either enhanced protein catabolism or impaired protein anabolism. In myoblasts exposed to DM-succinate, we did not observe an increase in mRNA transcripts encoding for E3 ubiquitin ligases (MuRF1, Atrogin-1) typically associated with enhanced muscle catabolism (**Fig. 1D**). We did, however, observe a reduction in protein anabolism as determined by an O-propargyl-puromycin (OPP) incorporation assay, performed in proliferating C2C12 myoblasts treated with 8 mM DM-succinate for 48 hours (**Fig. 1E**). Interestingly, differentiated C2C12 myotubes treated with 8 mM DM-succinate for 24 hours post myogenic differentiation did not exhibit statistically significant alterations in mean myotube diameter (**Fig. 1F**), suggesting that the DM-succinate-associated protein anabolism deficits are not sufficient to drive muscle atrophy.

### Elevated succinate levels impair *in vitro* myogenesis

To gain a more comprehensive understanding of the molecular changes associated with succinate elevation, we performed RNA sequencing (RNA-seq) on proliferating C2C12 myoblasts treated with 8 mM DM-succinate for 48 hours. Analysis of the transcripts involved in myogenesis and lineage specification revealed widespread transcript down-regulation after exposure to DM-succinate (**Fig. 2A**). These results led us to more closely investigate myogenic progression by differentiating C2C12 myoblast cultures in low serum conditions for 5 days in the presence or absence of 8 mM DM-succinate. In this assay, we assessed the degree of terminal differentiation by quantitative RT-PCR analysis of myogenic transcript expression (Pax7, MyoD1, Myog, and MyH1) and the number of nuclei fused into differentiated, Myosin Heavy Chain-positive myocytes/myotubes. Analysis of key myogenic transcript expression patterns supported the RNA-seq data, showing downregulation of MyoD1, Myog and MyH1, particularly earlier in the differentiation time course (**Fig. 2B**). Although not statistically significant, Pax7 expression was trending upwards in DM-succinate-treated cells (**Fig. 2B**). Differentiating C2C12 cells in the presence of 8 mM DM-succinate resulted in an impaired ability to form mature, multinucleated Myosin Heavy Chain-positive structures (control, 4 mM DM-succinate, 8 mM DM-succinate mean nuclei/myotube values = 5.43, 4.62, 2.98, respectively; **Fig. 2C, D**). Together, these data show that elevated succinate in muscle cells influences protein anabolism and suppresses myogenic differentiation and that these changes are distinct from direct myotube atrophy via enhanced protein catabolism.

**Figure 2:**
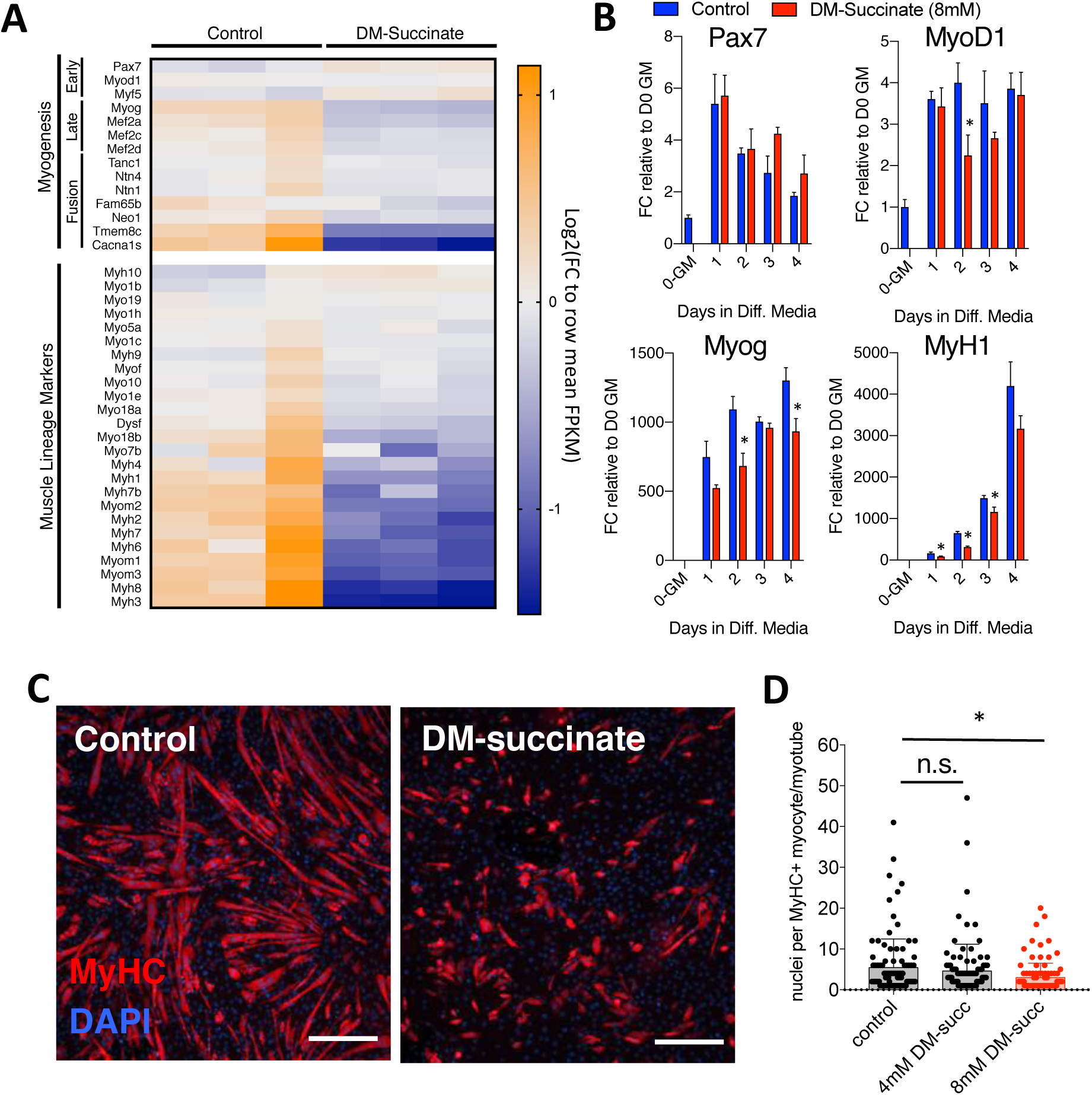
Elevated succinate levels impair *in vitro* myogenesis. **(A)** RNA sequencing analysis of proliferating C2C12 myoblasts treated with 8 mM DM-succinate for 48 hours. Heatmap depicting normalized FPKM values for myogenic and muscle lineage marker expression values. Blue=lower relative expression, orange=higher relative expression. n=3 replicates for each condition. **(B)** qPCR analysis of myogenic markers Pax7, MyoD1, Myog, and MyH1. Bar graphs depicting mRNA expression FC relative to D0-growth media at 0, 1, 2, 3, and 4d in differentiation media +/- 8 mM DM-succinate. n=3 replicates for each timepoint/condition, *p<0.05 between control and succinate samples for marked timepoints, as determined by multiple t tests with Holm-Sidak multiple testing correction. **(C)** Representative images of C2C12 cultures in the presence of differentiation media supplemented with 8mM DM-succinate for the entirety of differentiation (5 days). Red depicts myosin heavy chain positive myocytes and maturing myotubes. Nuclei are shown in blue (DAPI). Scale bar=100um. **(D)** A graph depicting the number of nuclei per myosin heavy chain positive structure (one point=one structure). Red=statistically significant (p<0.05) mean of biological replicates compared to control using unpaired, nonparametric, Mann-Whitney tests. n=4 replicates control/8mM DM-succinate, n=5 replicates 4mM DM-succinate, 20-40 myotubes/replicate.

### *In vivo* succinate supplementation impairs skeletal muscle homeostasis

Defects in progenitor cell activation, expansion, differentiation, or fusion compromise the regenerative capacity of skeletal muscle. Given our observations that DM-succinate impairs myogenic differentiation *in vitro*, we next asked if succinate supplementation negatively impacts *in vivo* skeletal muscle homeostasis. Wildtype mice were either supplemented with vehicle (1% sucrose) control drinking water or water with 2% succinate (+1% sucrose vehicle). We found that there was no significant difference in total body weight across the supplementation, or at the endpoint (**Fig. 3A, B**). In addition to total body weight, we longitudinally monitored body composition of the mice twice weekly for six weeks using echo magnetic resonance imaging (echoMRI). Total lean mass, as measured by echoMRI-based body composition analyses, did not show a difference between vehicle and succinate-supplemented mice (**Fig. 3C)**. Comparisons of *tibialis anterior* (TA) and *gastrocnemius* (GR) muscle wet weights after six weeks of supplementation did not reveal significant differences between experimental groups (**Fig. 3D)**. Following six weeks of succinate supplementation, we harvested TA muscles and stained tissue cross sections with laminin, a protein on the myofiber membrane, to evaluate myofiber size (**Fig. 3E)**. We quantified myofiber minimum feret diameters using Myovision^44^ and found a statistically significant (p<0.0001) decrease (∼6%) in the mean of non-linear regressions fit to the histograms of minimum feret diameter distribution between control and succinate supplemented cohorts (**Fig. 3F, Supplementary Fig S1A, B)**.

**Figure 3:**
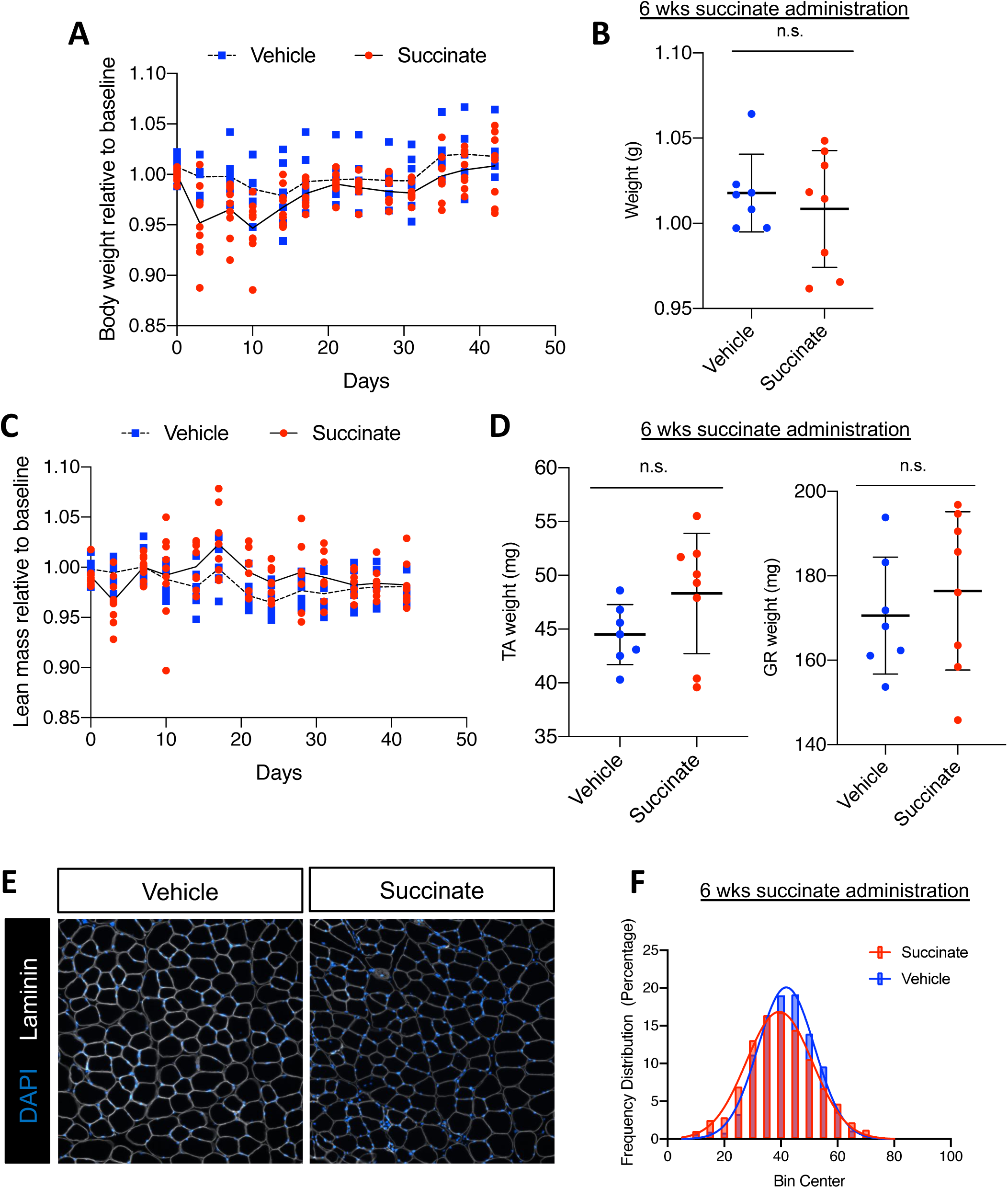
Succinate supplementation impairs skeletal muscle homeostasis. **(A)** Bi-weekly body weight tracking over 6-week succinate supplementation. Weights are normalized to pre-supplementation baseline measures. Individual points are individual animals, line represents the mean at each time point. **(B)** Mouse weight at endpoint, normalized to pre-supplementation baseline measures. **(C)** Bi-weekly echoMRI-measured lean mass normalized to pre-supplementation baseline measures. Individual points are individual animals, box represents the quartiles and whiskers represent the minimum and maximum values. **(D)** Weights of tibialis anterior (TA) and gastrocnemius (GR) at endpoint in milligrams. Individual points are individual animals, box represents the quartiles and whiskers represent the minimum and maximum values. **(E)** Representative images of TA muscle tissue sections from vehicle or succinate supplemented mice after 6 weeks administration. Cross sections are immunofluorescent stained with an antibody targeting laminin (white). Nuclei are labeled with DAPI (blue). Scale bars=100 µm **(F)** Quantification of minimum feret diameter of myofibers from vehicle and succinate supplemented mice. Feret diameters were binned to a histogram and fit with a non-linear regression (gaussian, least squares regression). Succinate exposed myofibers were significantly smaller; P< 0.0001 by extra sun-of-squares F test. n.s. = not significant by student’s t test, n=7 vehicle, n=8 succinate-supplemented FVB male mice, 4-5 months old at the start of the study.

### Succinate supplementation impairs skeletal muscle regeneration

Muscle progenitor cells are required for efficient muscle regeneration following serious injury^23-25^. In order to challenge resident satellite cells chronically exposed to elevated succinate, after six weeks of supplementation, we injured TA muscles with intramuscular barium chloride injection (1.2%). At 7 days and 28 days post injury (dpi) we harvested TA muscle for histological analysis. At 7 dpi, we observed statistically significant (p=0.0006) differences in myofiber regeneration as evidenced by ∼16% reduction in the mean of non-linear regression curves fit to the minimum feret diameter distribution (**Fig. 4A, B, Supplementary Figure S1C, D**). Muscle regeneration post barium chloride injury is typically completed four weeks post injury in healthy, adult mice. When we analyzed minimum feret diameters at 28 dpi, the difference in the mean of non-linear regression curves fit to the minimum feret diameter distribution became more pronounced (∼48% reduction, p<0.0001) (**Fig. 4C, D, Supplementary Figure S1E, F**). Injured TA muscle in succinate supplemented mice exhibited large areas of poorly regenerated tissue (**Fig. 4C**). To further investigate the impacts of chronic succinate supplementation on the satellite cell population we looked at markers of myogenic differentiation and proliferation. We saw no significant difference in the number of total (PAX7+) or proliferating (PAX7+KI67+) muscle progenitor cells in succinate-supplemented animals at 7- or 28 dpi, in concordance with *in vitro* proliferation data (**Fig. 4E, F; top**). At 7-days post injury there was no difference in the number of MYOG+ differentiating muscle progenitor cells between succinate- and vehicle-supplemented mice. However, we did see a significant accumulation of differentiating muscle progenitor cells (MYOG+) in succinate-supplemented animals at 28 dpi, indicative of stalled or incomplete myogenesis. In comparison, vehicle-supplemented mice had low numbers of MYOG+ cells at 28 dpi likely due to effective completion of myogenesis (**Fig. 4E; bottom**). Taken together, these data show that succinate supplementation alters progenitor cell dynamics resulting in impaired skeletal muscle homeostasis and diminished skeletal muscle regeneration.

**Figure 4:**
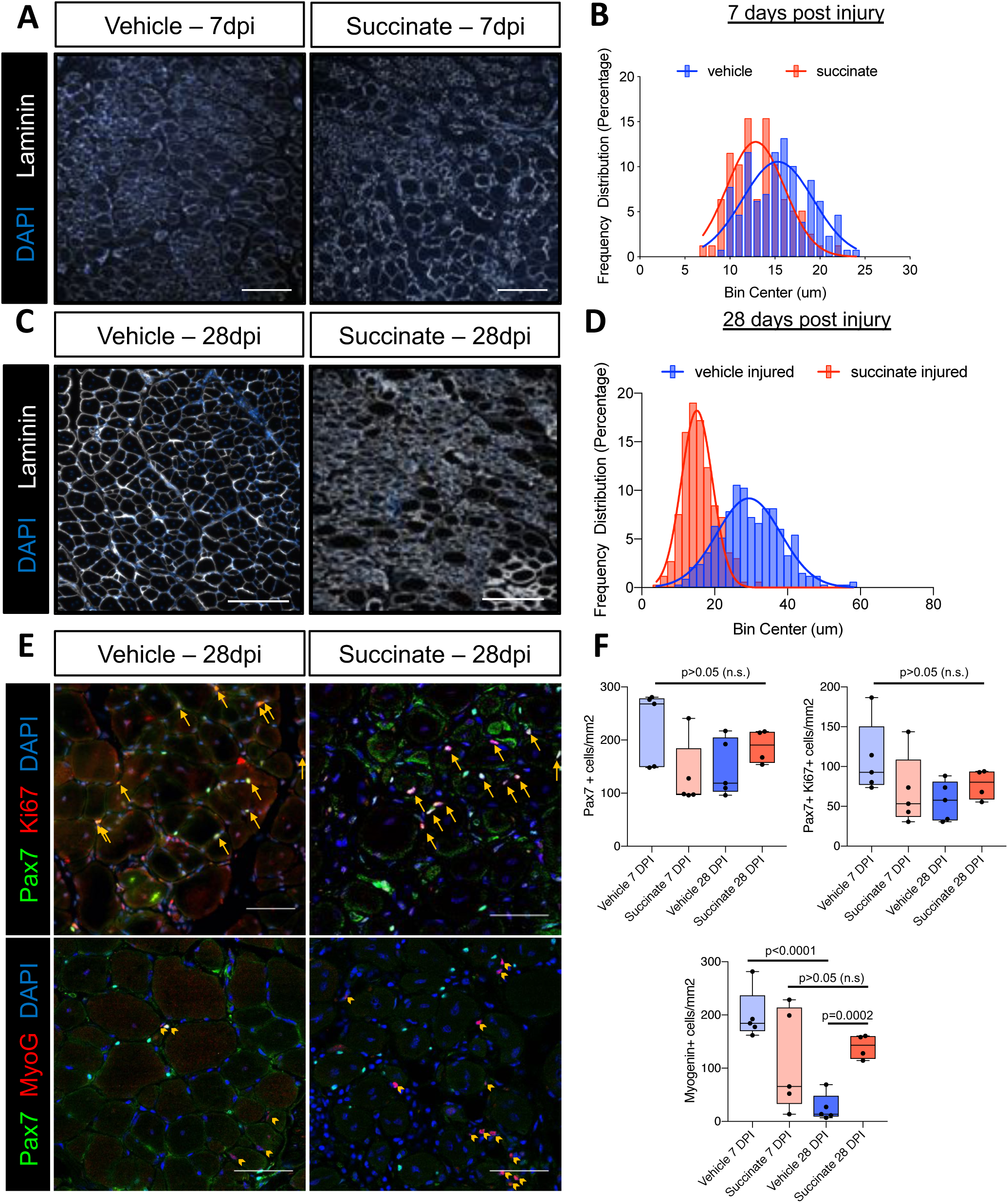
Succinate supplementation impairs skeletal muscle regeneration. **(A)** Representative images of TA muscle tissue sections from vehicle or succinate supplemented mice 7 days post 1.2% BaCl_2_ injury stained with an antibody targeting laminin (white). Nuclei are labeled with DAPI (blue). **(B)** A histogram of myofiber feret diameters at 7 days post injury (dpi). Non-linear regression and statistical analyses were performed as in Fig. 3, succinate injured myofibers were ∼16% smaller; p=0.0016. **(C)** Representative images of TA muscle tissue sections from vehicle or succinate supplemented mice 28 days post 1.2% BaCl_2_ injury stained with an antibody targeting laminin (white). Nuclei are labeled with DAPI (blue). **(D)** A histogram of myofiber feret diameters at 28 days post injury (dpi). Non-linear regression and statistical analyses were performed as in Fig. 3, succinate injured myofibers were ∼52% smaller; p<0.0001. **(E)** Representative images of TA muscle tissue sections from vehicle or succinate supplemented mice 28 dpi stained with antibodies targeting Pax7 (green), and Ki67 (red, upper panels), or Myogenin (red, lower panels). Nuclei are labeled with DAPI (blue). Yellow arrows in upper panels point to Pax7+/Ki67+ cells. Yellow arrow heads in lower panels point to Myog+ cells. **(F)** Quantification of Pax7+, Pax7+/Ki67+, and Myog+ cells per mm^2^. Data represent the quantification of a 0.844 mm^2^ area per animal. Data points represent individual animals. Box represents the inner quartiles and whiskers represent minimum and maximum values. p values are represented in the figure, as determined by unpaired student’s t test. Mice were 3-4 months old, FVB males. n=5 mice 28dpi vehicle; n=4 mice 28dpi succinate; n=5 mice 7dpi vehicle; n=3 mice 7dpi succinate. Scale bars in panel 4A and 4C=100 um, 4E=50 µm.

### Succinate elevation disrupts multiple metabolic networks in muscle cells

Succinate is gaining traction as a metabolite capable of acting as more than a substrate for succinate dehydrogenase and the respiratory chain^45^. Notably, succinate is located at the intersection of many diverse metabolic pathways and can influence a wide range of metabolic processes^46,47^. We performed non-targeted metabolomics profiling on control and 8 mM treated (48h) C2C12 myoblasts and found a striking number of differentially abundant metabolites (DAMs) between in the two conditions (214 features with p<0.05, FC>2). Network analyses of metabolic pathways associated with these DAMs implicated pathways involved in amino acid and nucleotide metabolism, as well as metabolic processes associated with mitochondrial metabolism and the TCA cycle (**Fig. 5A,B**). Considering the central role of the mitochondria in coordinating cellular metabolism and the known involvement of succinate in several mitochondria-coupled processes, we posited that succinate elevation could result in mitochondria dysfunction. Prior evidence suggests that myogenic differentiation involves metabolic rewiring characterized by increased mitochondria mass and a shift towards oxidative phosphorylation^48,49^. To assess mitochondria function, we measured cellular oxygen consumption rates (OCR) in proliferating myoblasts treated with 4 mM and 8 mM DM-succinate exposed to mitochondrial stressors. When treated with compounds targeting ATP synthase (oligomycin), the hydrogen ion gradient (FCCP), and complex I/III (Rotenone/Antimycin A), we noted altered OCR dynamics in myoblasts treated with the highest dose (8 mM) of DM-succinate (**Fig. 5C**). Maximal respiration, as determined by OCR measurements following the addition of the ionophore FCCP, was significantly reduced (∼35%) in the presence of 8 mM DM-succinate (**Fig. 5D**), thus significantly decreasing mitochondria reserve capacity^50^ (∼50% reduction; **Fig. 5E**). Basal respiration rates, and non-mitochondrial respiration were unaffected by succinate elevation (**Fig. 5F, G**). The ability of cells to respond to general mitochondrial stress was further assessed using an energy phenotyping test, which exposes cells to FCCP and oligomycin simultaneously and measures both ECAR (extracellular acidification rate) and OCR. When stressed, control cells exhibited expected increases in ECAR and OCR, whereas DM-succinate treated cells responded poorly (**Fig. 5H**). Taken together, these data show that while elevated succinate does not grossly affect mitochondria morphology or basal (resting) respiration parameters, it does impact the ability of cells to maximally engage mitochondrial respiration under stressful conditions.

**Figure 5:**
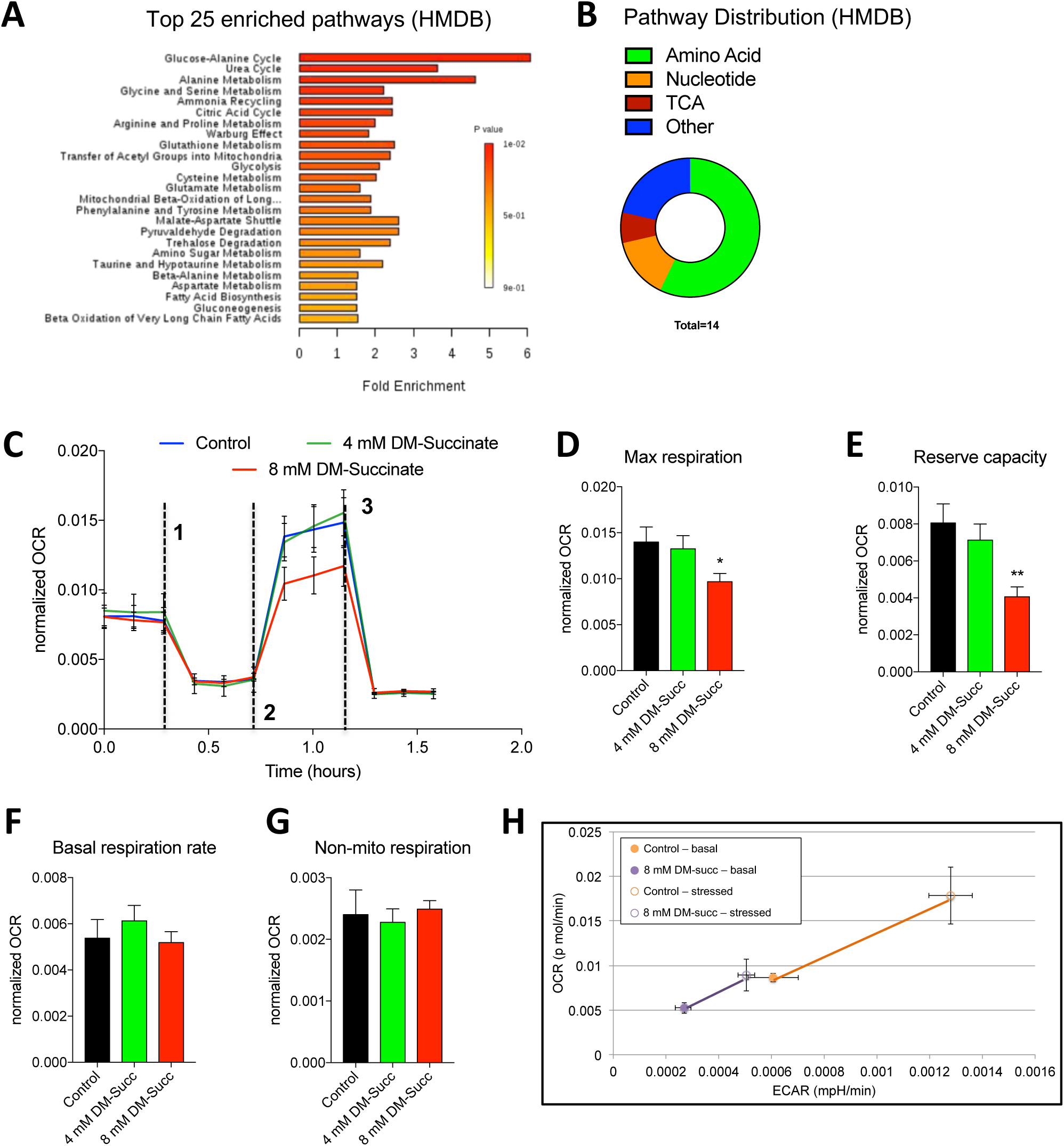
Succinate elevation impairs mitochondria function in myoblast cells. **(A)** A bar chart depicting pathway enrichment analyses (sorted by p-value) using HMDB IDs. Shown are the top 25 enriched processes. All data are derived from Metaboanalyst processing of non-targeted analysis of C2C12 cells treated with control or 8 mM DM-succinate media for 48 hours. n=3 control and n=3 8 mM DM-succinate. **(B)** A circular pie chart depicting the relative distribution of differentially regulated metabolic pathways using HMDB IDs during function explorer analysis. **(C)** Line graphs showing oxygen consumption rates (OCR), normalized to cell count, of C2C12 myoblasts treated sequentially with oligomycin (dashed line 1), FCCP (dashed line 2), and Antimycin A/Rotenone (dashed line 3) in the presence or absence of 4 mM or 8 mM DM-succinate. **(D-G)** Bar graphs displaying basal respiration rates (F), maximal respiration (G), reserve capacity (H), and non-mitochondrial respiration (I). n=3; *p<0.05, **p<0.01 by unpaired student’s t-test**. (H)** Line graphs depicting OCR and ECAR (extracellular acidification rate) in response to mitochondria stress (Energy phenotype assay; Seahorse) in C2C12 control or DM-succinate treated myoblasts. Filled circles are basal conditions, open circles are stressed conditions. n=3 control and n=3 8 mM succinate.

## DISCUSSION

Mitochondria and tricarboxylic acid (TCA) cycle defects are commonly observed in wasting skeletal muscle^51-53^. Direct alterations to TCA metabolite stoichiometry could therefore, in principal, contribute to functional deficits associated with muscle wasting. Indeed, multiple studies show that perturbations in the succinate/alpha-ketoglutarate ratio underlie a spectrum of cellular defects ranging from epigenome alterations to mitochondria deficits^15,19,54^. In this study, we found that succinate elevation in muscle progenitor cells impaired myogenic differentiation *in vitro* and disrupted *in vivo* muscle regeneration in response to injury. While this study did not directly assess the role of muscle progenitor cells in cancer-associated cachexia, we provide proof-of-concept evidence that tumor-associated factors (ie. succinate) are capable of functionally perturbing myogenesis, a process implicated by several groups as a contributing factor to muscle wasting^6,27,30,35,38^. We anticipate that future identification and evaluation of other tumor-associated metabolites will reveal many new drivers of muscle wasting and will prompt a re-evaluation of therapeutic strategies designed to limit cancer-associated weight loss.

Several groups show that elevated succinate can influence multiple metabolic processes including NADH/NAD+ redox state, phosphate metabolism, carbon source utilization, amino acid incorporation, and fatty acid synthesis^46,47,55,56^. Of note, succinate CoA ligase (SUCL) – which catalyzes succinyl-CoA and ADP/GDP to CoASH, succinate, and ATP/GTP – is subject to inhibition by the accumulation succinate^45^. This product inhibition of SUCL could lead to a buildup of upstream CoA species such as propionyl-CoA and methylmalonyl-CoA, which in turn would likely inhibit the catabolism of many macromolecules that are normally metabolized to succinyl-CoA, including branched-chain amino acids, cholesterol, propionate, and odd-chain fatty acids^57,58^. Future studies focusing on how succinate engages other metabolic networks will help to clarify how elevated succinate levels result in impaired muscle repair/regeneration.

Recent work shows that mitochondria and energy homeostasis play important roles in the regulation of myogenesis. A study that measured the relative contribution of oxidative phosphorylation (OXPHOS) to total cellular ATP in muscle cells demonstrates a shift from ∼30% OXPHOS dependent ATP production in proliferating myoblasts to ∼60% in terminally differentiated myotubes^59^. Accordingly, mRNA expression profiles and activities of mitochondrial enzymes including citrate synthase, isocitrate dehydrogenase, 3-hydroxyacyl-CoA dehydrogenase, cytochrome oxidase, NADH dehydrogenase, and succinate dehydrogenase are markedly increased during myogenic differentiation^60^. Several lines of evidence show that functional impairment of mitochondria negatively regulates myogenic differentiation. First, mitochondrial protein synthesis inhibition by chloramphenicol impaired chick embryo myoblast fusion, even when proliferation/apoptosis rates were maintained by tryptose phosphate broth or nucleoside supplementation^61^. Second, over-expression of key factors involved in mitochondria biogenesis (PGC-1 genes) in skeletal muscle enhanced mtDNA content, mitochondrial enzyme activities, and exercise performance^62^, whereas mice lacking PGC-1 show reduced mitochondria numbers, decreased expression of mitochondrial genes, and diminished muscle function^63,64^. We found that while DM-succinate impaired the maximal respiratory capacity of myoblasts, DM-succinate failed to affect basal mitochondria bioenergetics parameters or gross mitochondria morphology. Accordingly, proliferation and apoptosis rates were unchanged in DM-succinate treated cells. These data support the hypothesis that elevated succinate levels specifically impair myogenic differentiation, an energetically demanding process that challenges myoblast mitochondria.

Our data show that *in vivo* succinate supplementation impairs muscle morphometric parameters and regeneration properties, thus adding to the growing list of studies that report metabolite-driven skeletal muscle mass and/or functional alterations^65,66^. Cancer-associated lean mass loss is considered a metabolic disorder with complex and overlapping mechanisms of action (including both increased catabolism and impaired regeneration) that ultimately lead to a decline in skeletal muscle mass and function. While cytokines and chemokines have garnered the most attention as drivers of muscle wasting phenotypes, efforts to target, block or neutralize specific wasting factors have seen little success^67-69^. Our data underscore the importance of metabolites as regulators of muscle progenitor cell function, bringing attention to an understudied therapeutic avenue.

## ACKNOWLEDGEMENTS

The authors wish to thank members of the Doles lab for helpful discussions and manuscript suggestions and the Mayo Microscopy and Cell Analysis Core for experimental and technical support. J.D. was supported by the National Institutes of Health/National Institute of Arthritis, Musculoskeletal and Skin Diseases (NIH/NIAMS) R00AR66696, Mayo Clinic start-up funds, Career Development Awards from the Mayo Clinic SPORE in Pancreatic Cancer (NIH/ National Cancer Institute (NCI) CA102701) and the American Association for Cancer Research/Pancreatic Cancer Action Network, and the Glenn Foundation for Medical Research. P.C.A was supported by the Mayo Clinic Regenerative Sciences Training Program (RSTP). Metabolomics studies were made possible by the Mayo Clinic Metabolomics Resource Core through NIH/National Institute of Diabetes and Digestive and Kidney Disease (NIDDK) U24DK10049 originating from the NIH Director’s Common Fund. The authors of this manuscript certify that they comply with the ethical guidelines for authorship and publishing in the Journal of Cachexia, Sarcopenia and Muscle^70^.

## AUTHOR CONTRIBUTIONS

Study design: PCA, KAH, JDD. Data collection: PCA, KAH, AS, AMS. Data analysis/interpretation: PCA, KAH, JDD. Writing and editing of manuscript: PCA, KAH, AJ, JDD. All authors approved this manuscript.

