## Supplemental figure 1 for "The wasting-associated metabolite succinate disrupts myogenesis and impairs skeletal muscle regeneration"

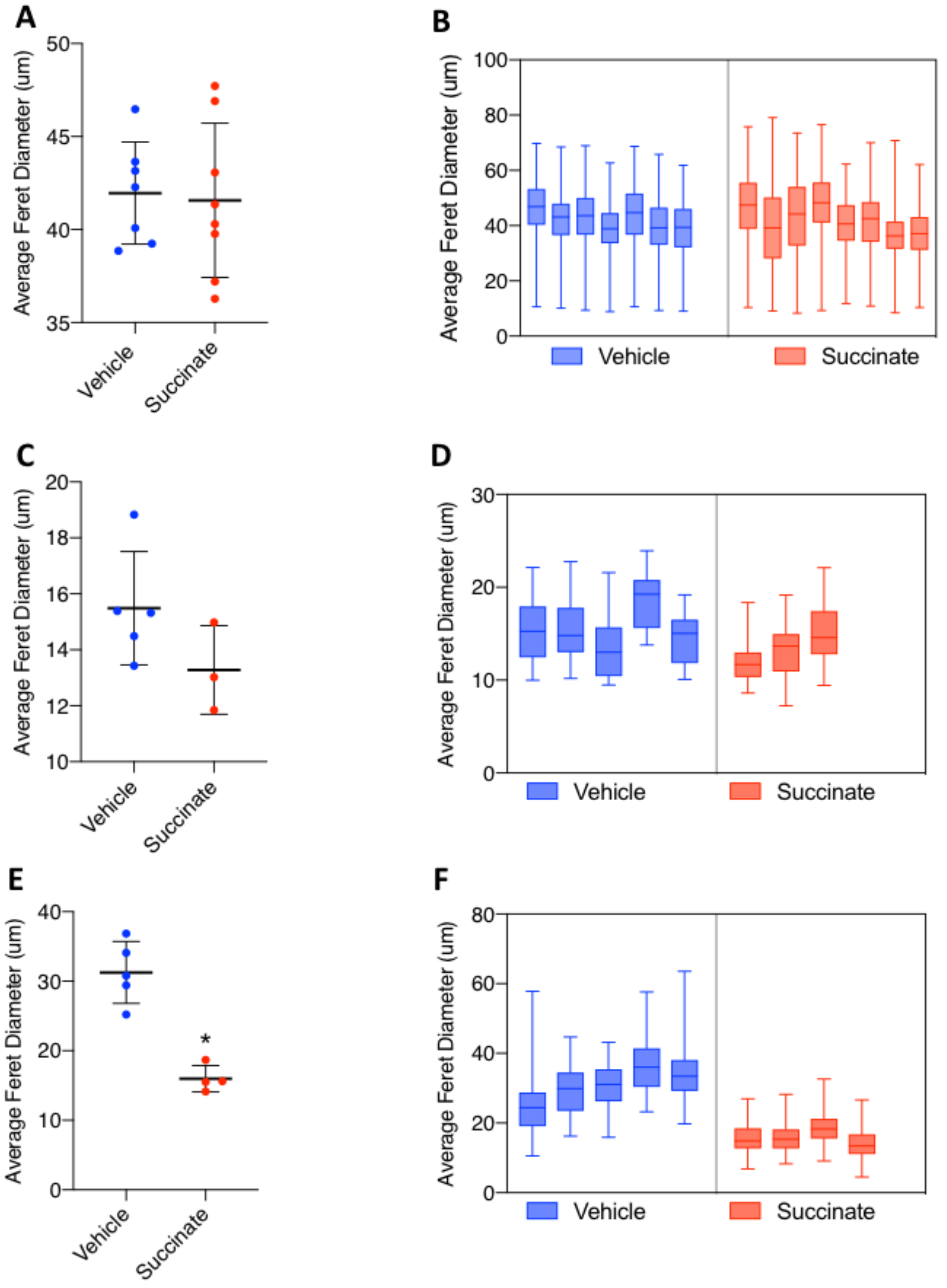

**Supplemental Figure 1: Individual assessment of myofiber regeneration (A)** Average feret diameter (um) of mice administered 6-week succinate supplementation (or sucrose vehicle). No significant difference between the average feret diameter (p=0.8374). Individual points are individual animals, line represents the mean, error bars are SD. n= 7 vehicle, 8 succinate. **(B)** Average feret diameter (um) and distribution for each individual mouse from panel A. Box represents the inner quartiles and whiskers represent minimum and maximum values. For A and B, total fibers quantified from all animals = 3,207 vehicle, 4,547 succinate. **(C)** Average feret diameter (um) of vehicle or succinate supplemented mice 7 days post 1.2% $BaCl_2$ injury. No significant difference between the average feret diameters (p=0.8374). Individual points are individual animals, line represents the mean, error bars are SD. n= 5 vehicle, 3 succinate. **(D)** Average feret diameter (um) and distribution for each individual mouse from panel C. Box represents the inner quartiles and whiskers represent minimum and maximum values. For C and D, total fibers quantified from all animals = 129 vehicle, 78 succinate. **(E)** Average feret diameter (um) of vehicle or succinate supplemented mice 28 days post 1.2% $BaCl_2$ injury. The average myofiber size was significantly smaller in succinate supplemented mice (p = 0.0004) Individual points are individual animals, line represents the mean, error bars are SD. n= 5 vehicle, 4 succinate. **(F)** Average feret diameter (um) and distribution for each individual mouse from panel E. Box represents the inner quartiles and whiskers represent minimum and maximum values. For E and F, total fibers quantified from all animals = 494 vehicle, 331 succinate. All p values calculated by students t test.
